# Divergent age-dependent conformational rearrangement within Aβ amyloid deposits in APP23, APPPS1, and App^NL-F^ mice

**DOI:** 10.1101/2023.10.24.563716

**Authors:** Farjana Parvin, Samuel Haglund, Bettina Wegenast-Braun, Mathias Jucker, Takashi Saito, Takaomi C Saido, K Peter R Nilsson, Per Nilsson, Sofie Nyström, Per Hammarström

## Abstract

Amyloid plaques composed of fibrils of misfolded Aβ peptides are pathological hallmarks of Alzheimer’s disease (AD). Aβ fibrils are polymorphic in their tertiary and quaternary molecular structures. This structural polymorphism may carry different pathologic potency and can putatively contribute to clinical phenotypes of AD. Therefore, mapping of structural polymorphism of Aβ fibrils is valuable to understand disease mechanisms. Here, we investigated how Aβ fibril morphology *in situ* differs in Aβ plaque of different mouse models expressing familial mutations in the AβPP gene. We used a combination of conformation-sensitive luminescent conjugated oligothiophene (LCO) ligands, Aβ-specific antibodies, and different fluorescence microscopy techniques. LCO fluorescence mapping revealed that mouse models APP23, APPPS1, and *App*^*NL-F*^ have different fibril structures depending on AβPP-processing genotype. Co-staining of Aβ-specific antibodies showed that individual plaques from APP23 mice, expressing Swedish mutations (NL) have two distinct fibril polymorph regions of core and corona. The plaque core is predominantly composed of compact Aβ40 fibrils and the corona region is dominated by diffusely packed Aβ40 fibrils. On the other hand, the APP knock-in mouse *App*^*NL-F*^, expressing Iberian mutation (F) along with Swedish mutation has tiny, cored plaques consisting mainly of compact Aβ42 fibrils, vastly different from APP23 even at elevated age up to 21 months. Age dependent polymorph maturation of plaque cores observed for APP23 and APPPS1 mice >12 months, was minuscule in *App*^*NL-F*^. These structural studies of amyloid plaques *in situ* can map disease-relevant fibril polymorph distributions to guide the design of diagnostic and therapeutic molecules.

**Significance:** Alzheimer’s disease (AD) is associated with the formation of deposits in the brain known as Aβ-amyloid plaques. AD can emerge as a sporadic disease or due to familial mutations in genes encoding for Aβ precursor and processing proteins. The Aβ-amyloid found in plaques displays different structures in sporadic AD and in various types of familial AD. We hypothesize that understanding plaque morphology and development is crucial for understanding the initiation and progression of AD. We here compared amyloid structures in three of the most used mouse models of human Aβ-plaque formation. Our findings suggest significant differences in plaque morphologies and structural maturation processes during aging. Our results emphasize that strain-like differences of Aβ-amyloids develop as a function of Aβ precursor protein-processing genetics and age.

## Introduction

Alzheimer’s disease (AD) is a neurodegenerative disease that affects millions of people worldwide. The manifestation of AD is complex and clinical signs span across cognitive, personality and behavioral changes, and motoric disturbances. Biochemical alterations, pathophysiological hallmarks, and neuroinflammation are obvious (1, 2). The major histopathologic findings in AD brain are Aβ-amyloid plaques (hereafter referred to as plaques), neurofibrillary tau tangles (hereafter referred to as tangles), and often cerebral amyloid angiopathy (CAA) from Aβ. Formation of plaques composed of Aβ peptides is tightly linked to the disease (3). However, this pathological hallmark of AD is also commonly found in healthy elderly (4), justifying the question of how plaque structures differ between healthy and diseased individuals and why. Although the link between plaque pathology and AD was first described by Alois Alzheimer in 1906 it is only during the later decades that a plethora of plaque morphotypes has been described systematically (5). Amyloid plaque and CAA microscopic morphology is likely associated with amyloid fibril structural polymorphism which is widespread for Aβ fibrils formed *in vitro* (6) and *in vivo* (7).

### Box 1

Benifits of conformation sensitive amyloid dyes

→ flexible backbones can **adapt to binding site**
→ flexibility renders differences in conjugation length generating **different optical output**
→ combinations of dyes give **increased contrast of structural variations** since different fibril polymorphs will have different binding sites with varying affinities
→ high **photostability** when bound to amyloid fibrils

Conformation sensitive amyloid ligands entail several benefits over conventional methods for staining amyloids (8) (9) (10) (11) (12) (13) (Box 1). Using two luminescent conjugated oligothiophenes (LCOs) qFTAA and hFTAA (14) in combination, we discovered that different polymorphs exist in plaque cores and periphery within the same plaque (8) in transgenic APPPS1 mice rich in Aβ42 (15). This difference was even more pronounced in APP23 mice, predominantly producing Aβ40 (16). For both mouse models we observed a change in plaque morphology and staining pattern as the mice aged (8).

Aβ-plaque polymorphism was also prolific when analyzing amyloid plaques in *postmortem* AD patient samples from familial (fAD) as well as sporadic AD using the LCO technology (17). This study strongly suggested that a cloud-like diversity of Aβ conformations appears within each patient. In addition, we recently demonstrated that there was a difference in plaque morphology between rapid progressing and slow progressing sporadic AD (18). Hence targeting specific polymorphs of Aβ aggregates is an attractive strategy for diagnostics and disease modifying therapies for ADs. Considering recently approved monoclonal antibody drugs targeting Aβ-amyloid (Aducanumab and Lecanemab) and patient specific response, it is important to understand Aβ turnover (19) and its plausible dependency on fibril polymorphism.

The first transgenic AD mouse model was introduced in the mid-1990ies (20) based on the current understanding of biochemical processing of the Amyloid-β precursor protein (AβPP) and how it is processed to form amyloid plaques. Since then, 197 mouse models of AD have been reported where of 77 are transgenic or knock-in for the AβPP gene and hence can be predicted to display Aβ plaque pathology (21) (22). Mouse models of AD will also in the future be crucial in the further search for disease relevant Aβ amyloid polymorphs.

The plaque forming Aβ peptide exists in several different isoforms, predominantly ending at amino acid 38 to 43. The peptides are formed by the cleavage of AβPP by several endogenous proteases according to the amyloidogenic cleavage pathway (3). Most mouse models of AD are based on a humanized AβPP gene flanked by familial mutations that promote the amyloidogenic processing pathway. The Swedish AβPP double mutation KM670/671NL (23) rendering overproduction of Aβ, is commonly used. Many mouse models are also combined with presenilin 1 (PS1) mutations to further exacerbate Aβ42 production.

In previous studies we found relatively low abundance of Aβ-amyloids displaying qFTAA fluorescence in APPPS1 mice compared to APP23 (8). This is likely a reflection of the lower qFTAA fluorescence we observed from *in vitro* formed recAβ1-42 fibrils compared to fibrils formed under the same conditions but from recAβ1-40 (8). This observation was confirmed by transmission electron microscope images (TEM) (24)(Fig. S1), which indicated less fibril bundling from Aβ1-42 than from Aβ1-40, suggesting a difference in fibril structure and the LCO binding pockets resulting from fibril bundling of the two different peptides (24).

It has been suggested that conformational variations, as reported by LCO staining, does not agree with the high-resolution studies of Cryo-EM structures (7), in that LCO staining show a wide variation of conformations in sAD (17) while Cryo-EM structures finds one predominant (type I) filament structure in sAD (7). Furthermore, fAD also had one predominant polymorph (type II) according to (7), where LCOs showed a separation, while still highly variable, depending on the type of fAD (17). Interestingly *App*^*NL-F*^and APP23 Aβ-amyloid filaments isolated and imaged by the same Cryo-EM procedure were reported to have the same main structure (type II) (7) (25). We therefore herein compared side by side Aβ-amyloid plaque conformational typing by LCO staining of three mouse models (Table S1, Fig. 1A).

**Figure 1:**
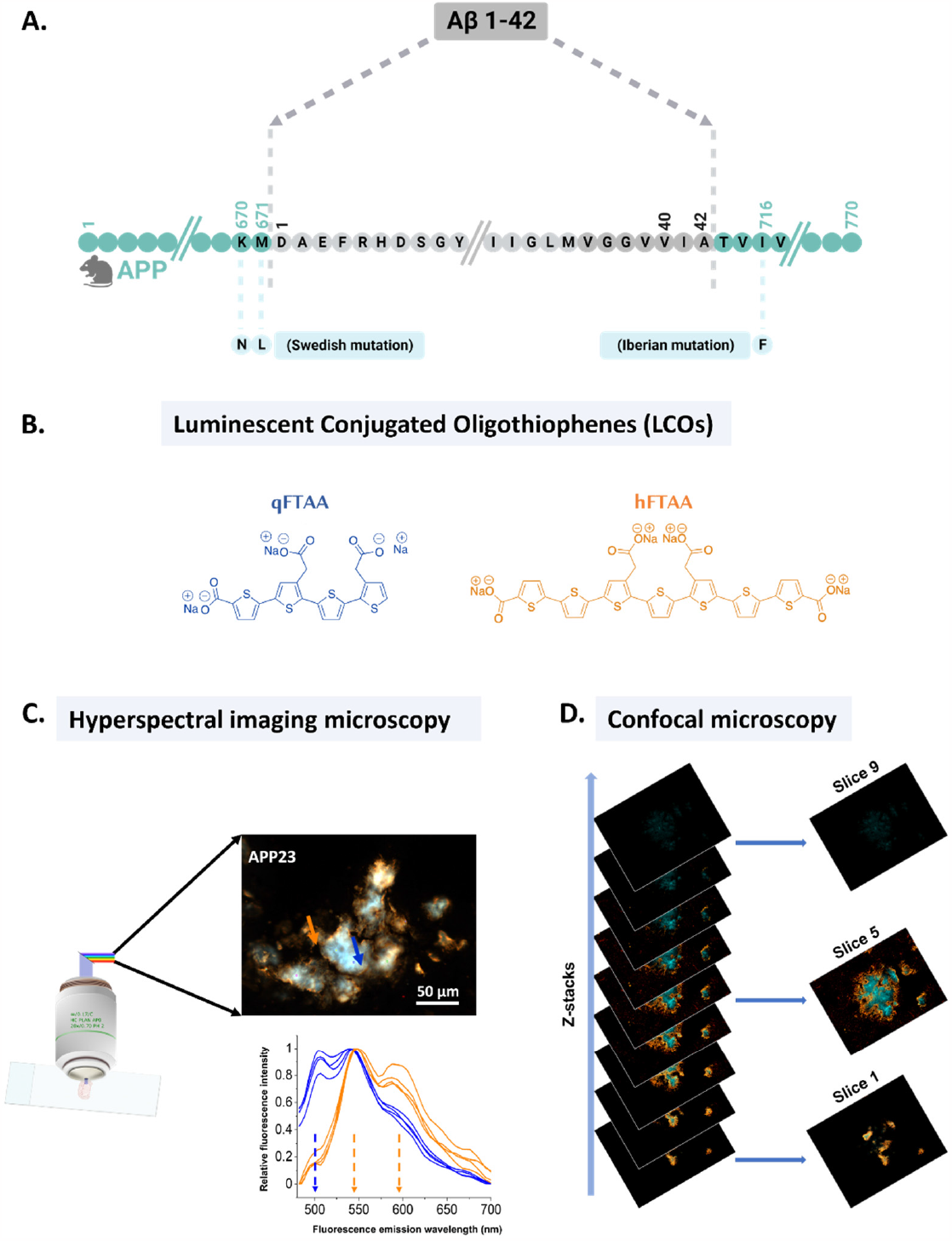
(A) Humanized mouse AβPP gene including Swedish mutation (KM670/671NL) for the APP23 mouse model mouse AβPP gene including the Iberian (I716F) mutation for the App^NL-F^ mouse model. (B) Thiophene-based conformation sensitive dyes: luminescent conjugated oligothiophenes (LCOs). (C) Hyperspectral fluorescence imaging of mouse brain section stained with LCOs: qFTAA and hFTAA. Both qFTAA and hFTAA give different emission spectra upon binding to amyloid fibril structures: qFTAA emits with a peak at 500 nm (blue arrow) meanwhile hFTAA emits with double peaks at 540 nm and 588 nm (orange arrows). (D) Z-stack confocal imaging of the mouse brain section provides information from different sections of the plaque.

The transgenic APP23 (16) has a 7-fold overexpression of human AβPP with the Swedish mutation (KM670/671NL) and produces more Aβ40 than Aβ42. The transgene is expressed under the Thy1 promoter element, resulting in production of AβPP mainly in neurons (26, 27). APPPS1 is a transgenic mouse model with three-fold overexpression of human AβPP with the Swedish mutation. In addition, it expresses a PS1 variant (L166P) that elevates the Aβ42/Aβ40 ratio. Also in this mouse, the transgene is expressed under the Thy1 promoter element (15). In the *App*^*NL-F*^ knock-in model mouse AβPP is expressed under the endogenous promoter ensuring physiological levels of AβPP at cell type and temporally relevant locations. The Aβ sequence was humanized and the insertion of the Swedish and the Iberian mutations (I716F) lead to a specific increase in Aβ42 production.

## Results and Discussion

We have for several years analyzed amyloid fibril deposits of different proteins and used thiophene-based ligands (28), and other molecular scaffolds such as trans-stilbenes (29) (30) for optical assignment of distinct protein aggregates *i*.*e*. amyloid fibril polymorphism on the folding and filament assembly levels (6). The notion behind this strategy is that different fibril polymorphs have different molecular structures of its ligand binding site (6) (Box 1). The two dyes, qFTAA (quadro-formylthiophene acetic acid) and hFTAA (hepta-formylthiophene acetic acid) (Fig. 1B), shows distinct spectral properties upon binding to different amyloid fibril structures (14). qFTAA upon binding to tightly packed mature fibril, fluoresces with an emission spectrum peaking at around 500 nm (24). hFTAA binds to both single filamentous and bundled fibrils, with red shifted emission spectra comprising peaks around 540 nm and 588 nm (24). It is known that Aβ amyloid plaque deposits have different microscopic morphologies when stained by immunohistochemistry and amyloid dyes. Aβ1-40 and Aβ1-42 amyloid fibril structural polymorphism is well documented by high resolution structural techniques of fibrils formed *in vitro* (31, 32) (33) (34) (35) in purified human (7) (36) and mouse brain (37) (25) derived amyloid fibrils, and in seeding experiments using brain derived fibrils as seeds for recombinant Aβ (38) (39). While the overall architecture is common with in-register parallel β-strands arranged in β-arches that comprise the filament structures; the fold, sequence arrangement of intermolecular interactions, and protofilament packing appear dramatically different in the Aβ fibril polymorphs (40). If and how the Aβ fibril polymorphs are associated with AD onset and progression is currently not established.

Co-staining of plaque from aged (18 Mo) APP23 mice, revealed two different fibrillar structural arrangements. Selected regions of interest (ROIs) from the core of the plaque were primarily occupied by qFTAA (blue-shifted spectrum with peak at 500 nm) indicating tightly packed mature fibrils (Fig. 1C - blue arrow). The core was surrounded by hFTAA stained ROIs (red-shifted spectrum with peaks at 540 and 588 nm) proposing immature fibrils in the periphery or corona (Fig. 1C - orange arrow). We supplemented the hyperspectral microscopy with confocal microscopy allowing use of multiple channels to utilize both the antibody and LCO staining at the same time, bringing out more detailed information about how the fibrils are organized in different parts of an individual plaque (Fig. 1D).

We then aimed for pairwise comparisons of Aβ-polymorphic differences between commonly used mouse models expressing human AβPP. The analysis affords resolution of the organization of structures allowed by optical microscopy (∼1 μm) but with the advantages of selective molecular probing with LCOs and observing intact amyloid structures in their near native environment using cryosections of flash-frozen brain (Fig. 2). We compared two AβPP-overexpressing transgenic mouse models (APP23 and APPPS1) but with different Aβ42/Aβ40 ratios (Table S1). We also compared AβPP knock-in model *App*^*NL- F*^ exhibiting endogenous AβPP-expression with a humanized AβPP transgene sequence with the over-expressor APP23. Using our established protocol for LCO discrimination of Aβ polymorphism as well as antibodies against different epitopes of the Aβ peptide to discriminate the two isoforms we deduced structural differences and how they corresponded to expression and dominating Aβ species.

**Figure 2:**
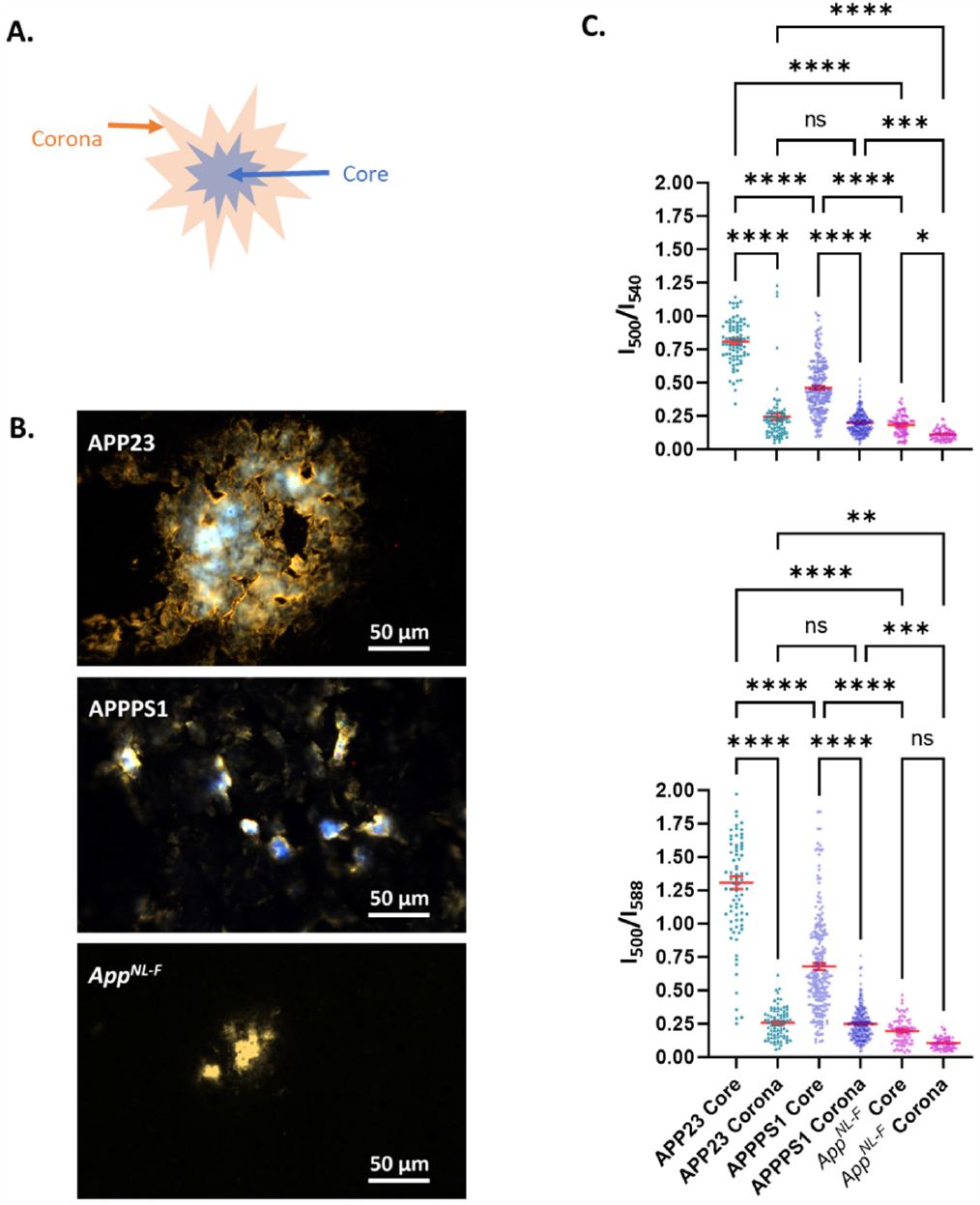
(A) Schematic representation of two distinct fibril polymorphic regions in plaque observed by double staining of qFTAA and hFTAA. The blue region represents the mature/bundled fibril-enriched plaque area termed as plaque core, dominated by qFTAA staining. The surrounding orange area of the plaque termed as corona, is enriched with hFTAA-stained diffusely packed fibrils. (B) Hyperspectral image overview of plaques stained with qFTAA and hFTAA from APP23, APPPS1 and App^NL-F^ mouse. The scale bars are 50 μm. (C) Fluorescence intensity ratiometric plot from the region of interest (ROI) from plaque cores and corona from APP23, APPPS1 and APP ^NL-F^ mouse. The error bars represent SEM. The upper panel shows the ratio of intensities at 500 nm and 540 nm, where 500 nm represents qFTAA emission and 540 nm represents hFTAA emission respectively. In the lower panel, the intensity of 540 nm is replaced by 588 nm, which represents another hFTAA emission peak. A pairwise one-way ANOVA test was performed for statistical analysis, where **** = p < 0.0001 and n.s. = non-significant.

Firstly, age-matched APP23 and APPPS1 mouse brain sections (18 months and 19 months respectively) were stained with a combination of qFTAA and hFTAA and full fluorescence spectra were collected using hyperspectral epifluorescence microscopy (41). 4 regions of interest (ROIs), each comprising 5×5 pixels (corresponding to ∼1×1 μm), from the core (Fig. 2A-B) and 4 ROIs from the corona (Fig. 2A-B) of the plaques were analyzed from each plaque. In total 15 images comprised 20 plaques for APP23 mouse, and 18 images contained 54 plaques for APPPS1 were analyzed. Fluorescence intensity ratiometric analyses were performed by division of the fluorescence intensity at 500 nm (qFTAA) with the fluorescence intensity at 540 nm and 588 nm (hFTAA) for each ROI (I_500_/I_540_ nm and I_500_/I_588_ nm). We first compared APP23 and APPPS1 mice. The analysis revealed a higher abundance of qFTAA fluorescence in the plaque cores of the APP23 model compared to APPPS1. In both mouse models the qFTAA fluorescence was higher in the core compared to the corona (Fig. 2C). This demonstrated that different transgenic genotypes have different fibril structure in the plaques depending on transgenic genotype and that the morphology differs between different parts (core & corona) of the same plaque (Fig. 2C).

Aβ fibrils generated *in vitro* render different packing architecture. Aβ1-42 fibrils are predominantly solitary while Aβ1-40 tend to form bundles comprising several fibril filaments (Fig. S1). Tightly packed Aβ-fibrils will render a higher qFTAA signal in *in vitro* experiments (24) and Aβ1-42 fibrils are to a lower degree than Aβ1-40 fibrils associated with high qFTAA fluorescence (8). Hence these results indicate that the fibrils are arranged differently in the core region (blue region in Fig. 2A) compared to the corona (orange region in Fig. 2A). It also indicates that the Aβ variant composition (Aβ40 vs Aβ42) can influence the plaque structure of the different mouse models. The data indicated that APP23 plaque core has more tightly packed fibrils compared to APPPS1 plaque core (Fig. 2C). The results were coherent with our previous data (8). Both these models overexpress Aβ but the Aβ42/Aβ40 ratio is different in that APP23 largely generates Aβ40 while APPPS1 has a 4.3-fold excess of Aβ42 (42)(Table S1). To delineate if total Aβ load or Aβ variant is the dominating denominator of polymorphic structure we compared the APP23 and APPPS1 mice with *App*^*NL-F*^ mice, again analyzing 19 plaques from 15 images of 18 months old mice. *App*^*NL-FF*^ mice is known to generate almost exclusively Aβ42 whereas AβPP expression are at endogenous levels (Table S1) (43). The *App*^*NL-F*^mice exhibited lower qFTAA fluorescence in both core and corona than the other two mouse models with the most striking difference between the three genotypes being observed in the plaque cores (Fig. 2C).

The analysis above was based on ocular determination of core and corona. Similar number of 5×5 pixel ROIs (∼1×1 μm) were collected from both core and corona regardless of size or number of plaques in each specific genotype. However, the size and number of plaques differ largely between the mouse models. APP23 mice exhibit large plaques as well as CAA (not analyzed here) while the plaques in APPPS1 mice are smaller and more abundant (Fig. 2B). The knock-in *App*^*NL-F*^mice, expressing endogenous amounts of AβPP present both fewer and smaller plaques than the overexpressing models, as can be expected (Fig. 2B).

To get an unbiased overall score of qFTAA versus hFTAA positivity we performed a whole image analysis that considers each pixel of the hyperspectral images collected. This analysis method will not report on the region-specific differences of plaque morphology but rather on the overall proportion and variability of amyloid staining within each image. This image analysis (described in Supporting methods and Fig. S2, Fig. S3) can be performed at any wavelength ratio and here we used I_500_/I_540_ nm and I_500_/I_588_ nm as in the ROI analysis. Our ROI based results showing that APP23 demonstrated the highest and *App*^*NL-F*^ mice the lowest abundance of qFTAA positivity (Fig. 2C) was confirmed in this unbiased image analysis (Fig. 3A-C, Fig. S3). APP23 showed considerably higher I_500_/I_540_ nm and I_500_/I_588_ nm ratios and a wide distribution compared to APPPS1. *App*^*NL-F*^ mice showed the lowest ratios and the narrowest distribution (Fig. 3A and Fig. S3E).

**Figure 3:**
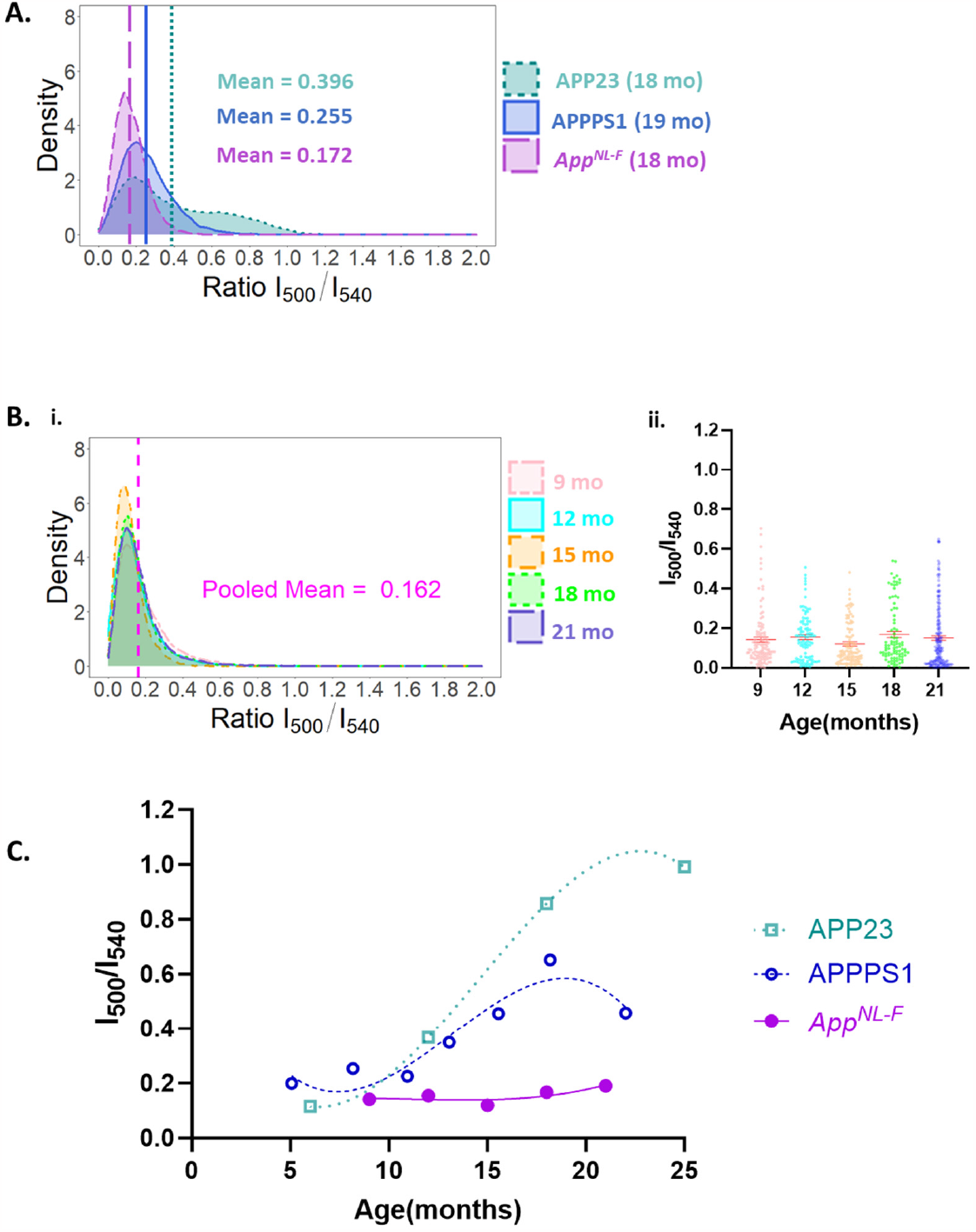
(A) Density plot of the APP23, APPPS1 and APP ^NL-F^ mouse showing the distribution of the calculated ratio matrix at I_500_/I_540_ nm. (B) The left panel (i) shows the density plot of App^NL-F^ mice at different ages. Total 31 images (32 plaques) for 9 months from 3 mice, 18 images (19 plaques) for 12 months from 1 mouse, 18 images (22 plaques) for 15 months from 1 mouse, 19 images (19 plaques) for 18 months from 1 mouse and 44 images (45 plaques) for 21 months aged from 3 mice were analysed. The pooled mean value represents the mean intensity ratio at I_500_/I_540_ nm of all the age groups of the App^NL-F^ mice. The right panel (ii) shows the intensity ratiometric plot from the collected ROIs from the core of the same mouse groups. The error bars represent SEM. (C) Diagram showing the comparison of the ratio of the intensity of the emitted light at 500 and 540 nm, I_500_/I_540_, of plaque cores versus mouse age of the App^NL-F^ mouse groups (filled circle, solid line) with APP23 (open rectangles, dotted line) and APPPS1 (open circle, dashed line). The lines are a third-order polynomial fitting of mean values of the intensity ratio versus age to show the trend. For APP23 raw data from 5 mice at 6 months, 7 mice at 12 months, 5 mice at 18 months and 5 mice at 25 months of age were analysed. APPPS1 raw data are compiled from a total of 19 mice (8). Note that the imaged Aβ-aggregates in 6 Mo old APP23 mice appeared to be intracellular inclusions and not bona fide Aβ-amyloid plaque cores.

Density plots were generated to visualize and compare the distribution of qFTAA/hFTAA I_500_/I_540_ nm fluorescence ratios between genotypes and within age groups of *App*^*NL-F*^ (Fig. S4A). This procedure was done for all three models at 18-19 months. The mean ratio for aged mice was 0.396 for APP23, 0.255 for APPPS1 and 0.172 for *App*^*NL-F*^ (Fig. 3A, Fig. S4). Interestingly the expression level of total Aβ is concomitant with an increasing proportion of Aβ40 in comparison with Aβ42 for models included in this study (Table S1). When analyzing the distribution of the qFTAA/hFTAA fluorescence ratio we find a trend in fluorescence signal tilting towards low qFTAA and high hFTAA positivity that corresponded both with decrease in AβPP expression levels and with increased Aβ42/Aβ40 ratio (Fig. 3A, Table S1).

*App*^*NL-F*^ mouse brain had a very low abundance of qFTAA positive plaques, also at 18 months. Differential qFTAA/hFTAA staining of plaque core versus corona developed during aging in both APPPS1 and APP23 mice, and increased qFTAA fluorescence of the plaque core as a function of mouse age with a transition at 12 months was reported by us as plaque core maturation (8). At very old age >18 months we previously observed a drop in plaque core qFTAA positivity due to increased hFTAA staining in APPPS1 mice (8). We therefore moved on to analyze the qFTAA/hFTAA ratio over several *App*^*NL-F*^ mouse ages, 9-21 months (Fig. 3Bii). We did not observe a noticeable trend of increased qFTAA fluorescence with *App*^*NL-F*^ mouse age (Fig. 3B, C). The I_500_/I_540_ nm ratio for the *App*^*NL-F*^mouse was around 3-fold lower compared to previously published aged matched APPPS1 mouse at 21 months (8) (Fig. 3C). The density plot of different age groups of *App*^*NL-F*^ mice at the intensity ratio matrix at I_500_/I_540_ nm also did not show any significant difference in the different ages (Fig. 3Bi), all the age groups are essentially overlapping (Fig. 3Bi). These results implied that there were not significant changes in the fibril structure over time. The individual density plot for each *App*^*NL F*^ mouse age group is represented in Supporting Fig. S4 side by side with the density plots of 18 months old APP23 mouse and 19 months old APPPS1 mouse.

Our previously published data on APPPS1 mice using the same method on the contrary showed a clear transition towards a more densely packed amyloid in the core of the plaques at ∼12 months, reflected by the increase in qFTAA fluorescence (Fig. 3C) and subsequent decrease >18 months (8), as discussed above. We therefore performed an analysis of APP23 plaque cores as a function of mouse age between 6 and 25 months to include in the comparison with *App*^*NL-F*^. APP23 plaque cores showed elevated I_500_/I_540_ nm ratios being even higher after 18 months than APPPS1 (Fig. 3C) making the discrepancy between *App*^*NL-F*^ mice and APP23 even stronger (Fig. 3C).

To further investigate the molecular basis for the Aβ-fibril polymorphism, co-staining with antibody and LCO was performed, and imaged by confocal microcopy. We here focused on the two mouse model extremes in qFTAA/hFTAA ratio and Aβ42/Aβ40 ratio, APP23 and *App*^*NL-F*^. The monoclonal antibody 4G8 (Aβ epitope 18-22) was used as a pan-Aβ detector while the antibody 12F4 (Aβ epitope 36-42) was used to selectively stain Aβ42 (Fig. 4A). The antibody-LCO co-staining revealed that APP23 plaque cores have qFTAA binding along with hFTAA and 4G8 antibody (Fig. 4B). The periphery or corona only showed hFTAA and 4G8 binding (Fig. 4B). 12F4 diffusely bound to the outermost part of the corona (Fig. 4B). This co-staining showed that individual APP23 plaque comprised two distinct fibril polymorph regions of core and corona. The plaque core consists predominantly of compact Aβ40 fibrils, meanwhile, the corona consists mainly of diffusely packed Aβ40 fibrils. Very little Aβ42 appeared to be present in APP23 Aβ-plaque as deduced from immunofluorescence (Fig. 4D). This result is coherent with previous mass spectrometry data for cored plaques in the similar mouse model APP_swe_ (44).

**Figure 4:**
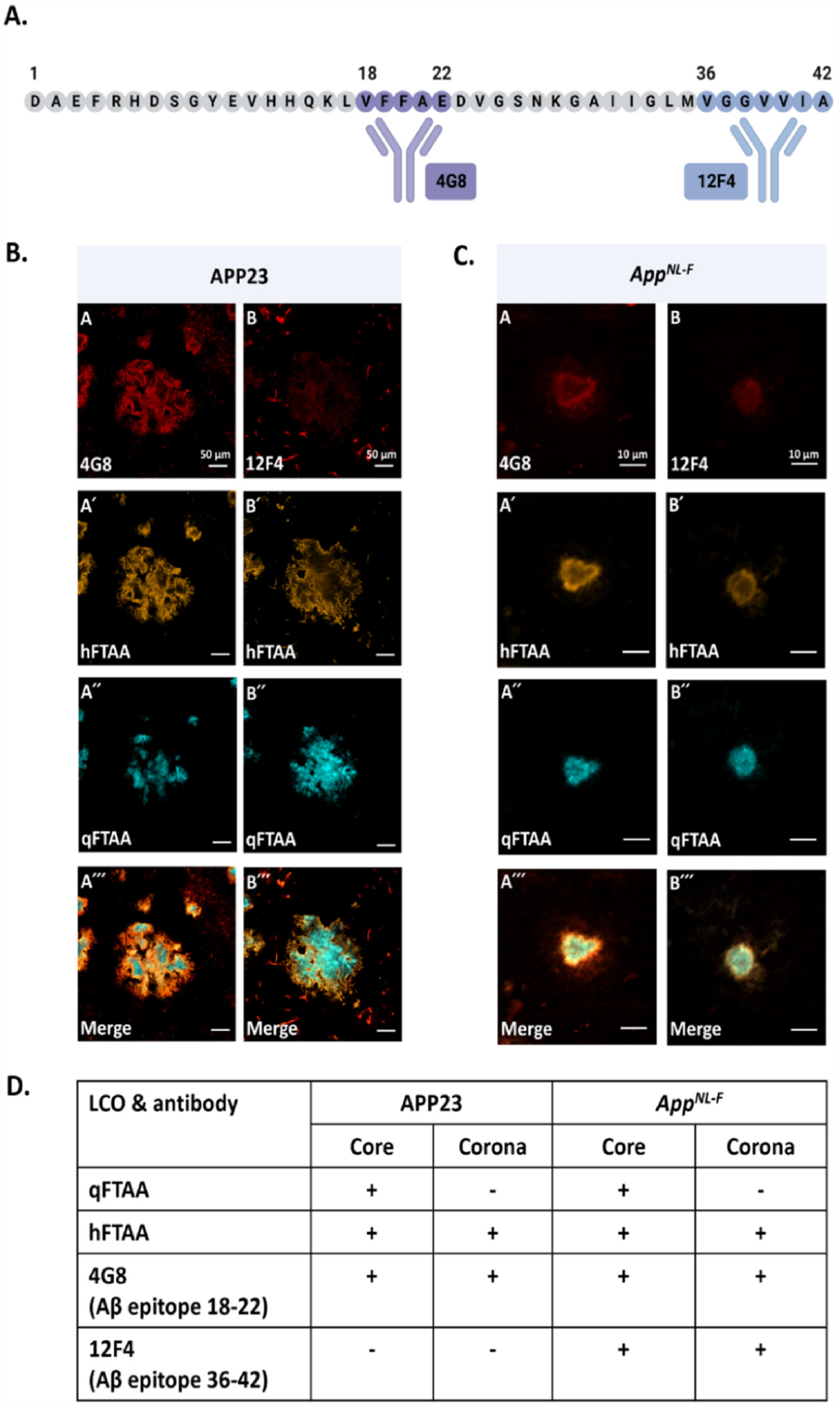
(A) Schematic representation of 4G8 and 12F4 antibodies. The 4G8 antibody recognizes the Aβ epitope sequence 18-22, meanwhile the 12F4 antibody recognizes the Aβ epitope sequence 36-42. (B) Antibody and LCO co-staining of plaque from APP23 mouse. Panel A shows 4G8 antibody staining, panel A’
s shows hFTAA staining, panel A″ shows qFTAA staining and panel A‴ shows the merged view of all three stains. Panel B shows 12F4 antibody staining, panel B′ shows hFTAA staining, panel B″ shows qFTAA staining and panel B‴ shows the merged view of all these three staining. The scale bars are 50 μm (C) Antibody and LCO co-staining of plaque from App^NL-F^ mouse. Panel A -A‴ and B-B‴ show similar antibody and LCO-co-staining as shown in (B). Note that the scale bars in (C) are 10 μm. Images in (B) and (C) are the single focal planes from the z-stack images in the confocal microscope where all the channels have maximum signal. (D) summary of LCO and antibody staining in different plaque regions in APP23 and App^NL-F^ respectively.

For *App*^*NL-F*^ mice, qFTAA positivity was not observable using epifluorescence hyperspectral microscopy (Fig. 2, Fig. 3). However, with confocal microscopy and optical sectioning we detected qFTAA positivity in tiny cores (Fig. 4C) of approximately 80 % of the plaques. These small cores are likely the reason for few areas with elevated I_500_/I_540_ nm ratios (Fig. 3Bii). Notably the cores do not significantly change with *App*^*NL-F*^ mouse age even with the unbiased image analysis approach (Fig. 3Bi). This demonstrates that the qFTAA positive part of each plaque is minute in comparison to >18 Mo APP23 where it is the dominating species. Co-staining with antibodies revealed that the plaque corona displayed hFTAA, 4G8 and 12F4 binding but no qFTAA fluorescence. Thus, we conclude that the qFTAA positive tiny plaque cores in *App*^*NL-F*^ is composed of compact Aβ42 fibrils, whereas most Aβ-plaques and the corona consist of diffusely packed Aβ42 fibrils. Hence it appears that the LCO staining preference predominantly reports on conformational differences rather than the ratio of Aβ42/Aβ40. The staining profiles of aged APP23 and *App*^*NL-F*^ plaque are summarized in Fig. 4D.

### Concluding remarks

The amyloid strain phenomenon and its dependence on fibril conformation has been extensively explored in the context of the prion protein and prion disease, where it is established that the prion structure correlates with disease phenotype (45). Amyloid fibril polymorphism coherent with what is known for prion strains appear evident for Aβ and AD (46, 47). The complexity of Aβ aggregation and the consequent heterogeneity of Aβ amyloid fibrillar structures was recently reviewed (48). Aβ-amyloid deposits as both plaques (40) and cerebral amyloid angiopathy (CAA) (49) and can be found in asymptomatic individuals. Although reaching old age many individuals never develop AD or other dementias. It is hence important to delineate the molecular details of the Aβ amyloid fibril polymorphs to better address the distribution of benign and disease relevant fibril types with molecular diagnostic and therapeutic strategies. The molecular tracers used *in vivo* in clinical practice today do not readily distinguish between disease related and non-disease related amyloid deposition (50). Furthermore, not all disease associated Aβ amyloids are detected by amyloid PET tracers. For example, carriers of the Arctic mutation (E22G), despite extensive Aβ fibril load, do not retain PiB in PET imaging (51). A wide distribution of LCO staining patterns in different patients suggest that the polymorphic patterns of different plaques are very hard to predict (17).

In this work we aimed to further understand the influence of isoform and expression level of the Aβ peptides on amyloid fibril polymorphism in different AβPP mouse models by direct staining and imaging of Aβ plaque *in situ*. We found that Aβ plaques in the three mouse models included in this study exhibit several different morphologies. The Aβ plaque morphology changes over time in APP23 and APPPS1 mice (8) but not in *App*^*NL-F*^ mice. *App*^*NL-F*^ mice contain small plaque dominated by hFTAA fluorescence. Plaque formation onset has been reported at 6 weeks of age in APPPS1 mice (15) and at 6 months of age in both APP23 (16) and *App*^*NL-F*^ (43) mice. This indicates that the age of the mouse or the age of the plaque cannot exclusively explain the difference in plaque morphology between APP23 and *App*^*NL-F*^ mice. Notably, *App*^*NL-F*^ and APP23 Aβ-amyloid filaments isolated and imaged by Cryo-EM were reported to have the same main structure polymorph (type II) (7) (25). To understand this discrepancy, it is conceivable that LCO staining reports on higher order assemblies of filaments with the same filament fold, or that Cryo-EM preparation (solubilized with sarkosyl, and centrifugation) and selective particle imaging influences the Cryo-EM results. The latter effect has recently been discussed in *in situ* Cryo-EM tomography of *App*^*NL-G-F*^ mice compared with Cryo-EM structure determination of isolated *ex vivo* fibril material (52).

Several PS1 mutations found in human fAD act by increasing the release of Aβ peptides from AβPP. However, the Aβ isoform varies between mutations. PS1-A431E results in high abundance of Aβ peptides but a low Aβ42/Aβ40 ratio (approximately 1/7^th^ for the mutant carrier compared to the average for cases of sporadic AD in the same study) (53). On the contrary carriers of PS1-E280A in a different study had an almost doubled Aβ42/Aβ40 ratio compared to sporadic AD cases (54) (both studies used ELISA to deduce the Aβ42/Aβ40 ratio). In other words, carriers of PS1-A431E generate more Aβ40 and PS1-E280A carriers generate more Aβ42 than do patients without mutation. In this sense, APP23 mice are like PS1-A431E while *App*^*NL-F*^ resemble PS1-E280A in terms of dominating aggregated Aβ peptide isoform. In studies of human AD cases, it has previously been shown that Aβ plaques in PS1-E280A carriers display very low qFTAA fluorescence (low intensity at 500 nm) while PS1-A431E carriers generate plaques with high qFTAA signature (17). This is in line with the LCO signatures of aged APP23 and *App*^*NL-F*^ mice described here. We hence conclude that time is an important factor for generation of the tightly packed cored plaques seen in APP23 mice and that the mature cored structure is promoted by the presence of abundant Aβ40. Plaque core maturation towards high qFTAA fluorescence is a feature much less pronounced in *App*^*NL-F*^ mice forming almost exclusively Aβ42 plaque. Hypothetically if *App*^*NL-F*^ mice were to be aged for a very long time >30 Mo it is conceivable that the qFTAA signature would increase. Also, the expression levels generating more Aβ in APP23 and APPPS1 mice than in *App*^*NL-F*^ mice may matter. However, this is not the full explanation. A previous study treating APPPS1 mice with a BACE-1 inhibitor that decreased total Aβ production and hence the number and size of Aβ-amyloid plaque in young mice, but the qFTAA/hFTAA ratio was altered by increasing in cortical regions and decreasing in thalamus, hypothalamus, and hindbrain regions (55). No difference was seen in treated versus non-treated 14 Mo old APPPS1 mice. These results indicate that the issue is more complex than merely Aβ-concentration. Our studies, considering the variable human pathology, strengthen the argument for translational work on using various mouse models as valuable prototypes for mapping Aβ-fibril polymorphism. The LCO-hyperspectral approach allows mapping the spatial distribution and the substructural organization of nearly intact amyloid structures *in situ* in their near native environment of formation.

### Methods in brief

*(Detailed methods are found in Supporting information)*

#### Preparation of tissue sections

Flash-frozen mouse brains of transgenic APP23 and APPPS1 and knock-in *App*^*NL-F*^ were used for making brain cryosections of 10 μm. Tissue sections were incubated with the LCO staining solutions comprising 2:1 qFTAA and hFTAA. After rinsing in PBS the tissue sections were air-dried and mounted with DAKO fluorescence mounting medium and cover slip.

For co-staining of antibodies and LCOs, Fixed and blocked tissue sections were then incubated with 4G8 and 12F4 primary antibodies respectively for overnight at 4°C. Following washing steps, tissue sections were incubated with Alexa Fluoro 594 secondary antibody. Then tissue sections were then incubated with LCOs and mounted as described above.

#### Hyperspectral Fluorescence Microscopy

Hyperspectral imaging was performed with a LEICA DM6000 B epifluorescence microscope equipped with a hyperspectral camera (ASI). A 436/10 nm excitation and 475 nm long pass emission filter was used for image acquisition.

From each image 4 regions of interest (ROI) were selected from the core and 4 ROI were selected from the corona of each plaque. Fluorescence intensity at 500 nm and 540 nm or 588 nm of the spectra from each ROIs were used to generate the intensity ratiometric plots I_500_/I_540_ or I_500_/I_588_ respectively. Representative images from each genotype were used for side-by-side analysis and comparisons in Fig. 2 and Fig. 3.

#### Confocal microscopy

Zeiss LSM780 confocal microscope was used to acquire z-stack images of LCO and antibody co-stained tissue sections. Argon 458, 488, 514 nm laser lines and DPSS 561-10 laser lines were used to excite the LCOs and Alexa Fluoro 594. Images were acquired with 20x objective.

#### Unbiased image analysis

The hyperspectral microscope Spectraview software saves one text file for the emission intensity for each pixel in by emission wavelength. Using an in-house generated program in RStudio, the ratio between intensity at 500 nm and 540 nm in each pixel was calculated. Pixels containing only background fluorescence were filtered out by the program (Fig. S3A-D). Violin plots and density distribution curves for the emission ratios for all images were generated (Fig. 3A,B and Fig. S3E).

## Supporting information

Parvin et al. Supporting information

## Acknowledgment

Microscopes were made available through ProLinC core facility for biophysical measurements at Linköping University.

FP is enrolled in the Forum Scientium graduate school at Linköping University.

