## Supplementary material for "Divergent age-dependent conformational rearrangement within Aβ amyloid deposits in APP23, APPPS1, and App^NL-F^ mice": Parvin et al. Supporting information

#### Supporting information contains:

Supporting methods

Supporting tables

Table S1. Overview of the Alzheimer's disease mouse models used in the study.

Table S2. A guideline to choose the values for filter setting for mouse tissue.

Supporting figures

Fig. S1 Bundling of *in vitro* generated Aβ1-40 fibrils.

Fig. S2 Filtration settings of LCO fluorescence data using a relative filter setting in RStudio.

Fig. S3 Generation of filtered LCO data for violin plots and density plots in RStudio.

Fig. S4 Density distributions of qFTAA/hFTAA at different ages of $App^{NL-F}$ mice.

#### Supporting methods

**Fibrillation of recombinant Aβ peptides and electron microscopy:** Fibrillation and electron microscopy of recombinant Aβ1-40 and Aβ1-42 peptides were performed as in (1). In short, the peptides were purchased from rPeptide, dissolved and stored as stocks at -20 °C in 2 mM NaOH at a concentration of 1 mg/ml. At time of fibrillation the peptides were diluted to a concentration of 10 µM in PBS and fibrillated at 37 °C without shaking. Samples were collected at the beginning of the equilibrium phase for fibril formation as deduced by ThT fluorescence. Carbon-coated copper grids were used to prepare TEM samples of fibrils negative stained with uranyl acetate. TEM images were collected using a Jeol JEM 1230 microscope at 100 kV using a Gatan CCD camera.

**Animals:** All animal experiments were conducted in agreement with protocols approved by the local Animal Care and Use Committees respectively. $App^{NL\text{-}F}$ mice were reared by Takashi Saito and Takaomi Saido lab at RIKEN Center for Brain Science, Tokyo, Japan. APP23 and APPPS1 mice were reared at Mathias Jucker lab at Hertie Institute for Clinical Brain Research, Tübingen, Germany. APP23 mouse tissues were handled as described previously (2). APPPS1 mouse tissues used here are described previously (3). This study hence allowed direct comparison of $App^{NL\text{-}F}$ with previous data of APP23 and APPPS1. Data from a total of 50 brains were included in the study: APPPS1 (n=19), APP23 (n=22), and $App^{NL\text{-}F}$ (n=9).

**Staining solutions:** The LCOs (qFTAA and hFTAA) were synthesized as described earlier (4). LCOs were dissolved in 2 mM NaOH in $dH_2O$ to have a stock solution of 1 mg/ml which gives a qFTAA solution of 1.8 mM and hFTAA solution of 1.1 mM. LCOs are stored at 4°C until further use. For double staining of mouse brain sections with LCOs, qFTAA was diluted to 1:10,000 and hFTAA 1:1392 and mixed in a ratio of 2:1 correspondingly. This gives a final staining solution of 120 nM qFTAA and 262 nM hFTAA. For antibody staining, 4G8 (Aβ epitope 18-22) and 12F4 (Aβ epitope 36-42) antibodies were diluted to 1:300 to stain mouse brain sections followed by Alexa Fluoro 594 diluted to 1:400.

**Preparation of tissue sections:** Flash-frozen 18 months old mouse brains of transgenic APP23 and APPPS1 and knock-in $App^{NL\text{-}F}$ were used for making brain cryosections of 10 µm. Cryosections were fixed in two consecutive ethanol concentrations of 96% (v/v) and 70% (v/v), 10 minutes for each concentration at room temperature. Tissue sections were further rehydrated with $dH_2O$ and PBS, pH 7.4, each step having 10 minutes of incubation time. Following the rehydration steps, tissue sections were incubated with LCOs (2:1 qFTAA and hFTAA) for 30 minutes. After incubation with LCOs, tissue sections were washed with PBS 3 times and incubated in PBS for 5 minutes. Later tissue sections were dried and mounted in DAKO fluorescence mounting medium.

For co-staining of antibodies and LCOs, the flash-frozen tissue sections were fixed at 70% (v/v) ethanol at 4°C for 3min. Prior to 70% (v/V) ethanol incubation, tissue sections were kept at room temperature for 30 minutes. Following ethanol incubation, tissue sections were rehydrated in $dH_2O$ for 2x2 minutes and in PBS for 10 minutes. Later tissue sections were blocked with 5% goat serum in PBS-T (0.1% triton x-100) at room temperature for 1 hour. Tissue sections were then incubated with 4G8 and 12F4 primary antibodies overnight at 4°C.

After primary antibody incubation tissue sections were washed with PBS-T for 3x10 minutes. Following the washing steps, tissue sections were incubated with Alexa Fluoro 594 secondary antibody at room temperature for 1 hour. Later tissue sections were washed with PBS for 3x10 minutes. Then tissue sections were incubated with LCOs (2:1 qFTAA and hFTAA) for 30 minutes. Following 30 minutes of incubation with LCOs, tissue sections were washed 3 times with PBS. Then tissue sections were dried well and mounted with DAKO fluorescence mounting medium.

**Hyperspectral Fluorescence Microscopy:** For Hyperspectral imaging of mouse brain sections, LEICA DM6000 B microscope was used, equipped with a spectral camera. A 436 nm long pass excitation filter (436/10 (LP475)) was used for image acquisition. Images were acquired with 20x objective. Images were acquired with 20x objective except for the mouse groups of $App^{NL-F}$ at different ages (9 months, 12 months, 15 months, 18 months and 21 months). These images were captured with 40x objective. The age series of $App^{NL-F}$ mice comprised a total of 9 mice (n= 3 at 9 Mo, n=1 at 12 Mo, n=1 at 15 Mo, n=1 at 18 Mo and n=3 at 21 Mo). For the age comparison of APP23 we analyzed a total of 22 mice (n=5 at 6 Mo, n= 7 at 12 Mo, n= 5 at 18 Mo, and n= 5 at 25 Mo). At 6 Mo mostly intracellular inclusions were present in APP23. For age comparison of APPPS1 data were plotted from (3), comprising a total of 19 mice.

Representative images from each genotype were analyzed. From each image, 4 regions of interest (ROI) were selected from the core and 4 ROIs were selected from the corona of each plaque. Fluorescence intensity at 500 nm and 540 nm of the spectra from each ROI were used to generate the qFTAA/ hFTAA ($I_{500}/I_{540}$) plot.

**Confocal microscopy:** Zeiss LSM780 confocal microscope was used to acquire z-stack image of LCO and antibody costained tissue sections. Argon 458, 488, 514 nm laser lines and DPSS 561-10 laser lines were used to excite the LCOs and Alexa Fluoro 594. Images were acquired with Plan-Apochromat 20x/0.8 M27 objective with a frame size of 1024x1024 and scanning area at zoom 1.0.

**Unbiased image analysis in RStudio:** Text files with intensity data at 500 nm, 540 nm and 588 nm for each image were loaded into RStudio as matrices. Relative filter settings were applied to remove unwanted signals, for example the dark background. The low wavelength matrix is used to filter out the high intensity noise because high intensities in the low wavelength matrix reflects bright noise. On the other hand, the high wavelength matrix is used to filter out the dark background noise. Filtering out the unwanted pixels should be done carefully with a robust reference interval to have reproducibility. Gaussian normal distribution of the pixels would be a suitable reference interval for that (Fig. S1). Calculating the mean value (μ) and using certain values of standard deviation (σ) from the mean value, the upper cut-off limit and lower cut-off limit can be set to the low wavelength (500 nm) and high wavelength matrices (540 nm). After several trial and error, certain upper and lower cut-off limits have been found to produce reproducible results for mouse brain tissue. These upper & lower cut-off limits differ for different tissue types, for example mouse brain tissue and fly brain tissue (*Drosophila melanogaster*). Even for mouse brain tissue, different mouse genotypes require different upper & lower cut-off limits to produce distinguishable results for different genotypes. The general guidelines have been mentioned in Table S2 describing how to set the cut-off limits for different genotypes based on the image outlook. After applying the filter setting, the ratio between the low wavelength matrix and high wavelength matrix is calculated and stored in a matrix. This new calculated ratio matrix is further used to illustrate data. A heatmap plot is generated to justify the effectiveness of the filter setting (Fig. S2). Moreover, the ratio matrix is used to generate a violin plot, like the ratiometric plot of ROI, but now on a larger full image scale (Fig. S3E).

### Supporting tables

***Table S1.*** *Overview of the Alzheimer's disease mouse models used in the study*

| **Mouse models/strains** | **APP23** | **APPPS1** | ***App^NL-F*** |
|---|---|---|---|
| Generation | First | First | Second |
| **Genetic modification** | **Transgenic** | **Transgenic** | **Knock-in (KI)** |
| **Promoter** | **mouse Thy1 element** | **mouse Thy1 element** | **mouse endogenous AβPP** |
| Humanized gene(s) | huAβPP$^{751}$(Swe) | huAβPP$^{695}$(Swe); huPSEN1(L166P) | huAβPP$^{751}$(Swe) |
| Mutations | AβPP $^{KM670/671NL}$ | AβPP $^{KM670/671NL;}$ PS1$^{L166P}$ | AβPP $^{KM670/671NL,I716F}$ |
| **AβPP expression level** | **7-fold overexpression** | **3-fold overexpression** | **1-fold expression** |
| **Ratio of Aβ amyloid (Aβ42/Aβ40)** | **0.2-0.42*** | **2.8-4.3*** | **11-8600**** |
| Availability | The Jackson Laboratory #stock 030504 | Mathias Jucker & The Jackson Laboratory #stock 3765351 | Takaomi Saido |
| Reference | Sturchler-Pierrat, C., et al., Proc Natl Acad Sci U S A, 1997. 94(24): p. 13287-92. | Radde, R., et al., EMBO Rep., 2006. 7(9): p. 940-6. | Saito, T., et al., Nat Neurosci, 2014. 17(5): p. 661-3. |

* 6 - 25 months for APP23
6 - 18 months for APPPS1

* * 6 - 21 months for *App^NL-F*

Ye, L., et al., EMBO Rep., 2017.18(9): p. 1536-1544.

**Table S2:** A guideline to choose the values for filter setting for mouse tissue.

| Outlook of the image | Lower limit | Upper limit |
|---|---|---|
| When the background is completely dark, the tissue does not show any fluorescence. More yellow fluorescence appears in the image than blue fluorescence. | $\mu$ | infinity ($\mu$+1000$\sigma$) |
| When the background is not completely dark but blurred with another color. The tissue shows intense blue fluorescence. | $\mu$+1$\sigma$ | infinity ($\mu$+1000$\sigma$) |

#### Supporting figures

A

B

C

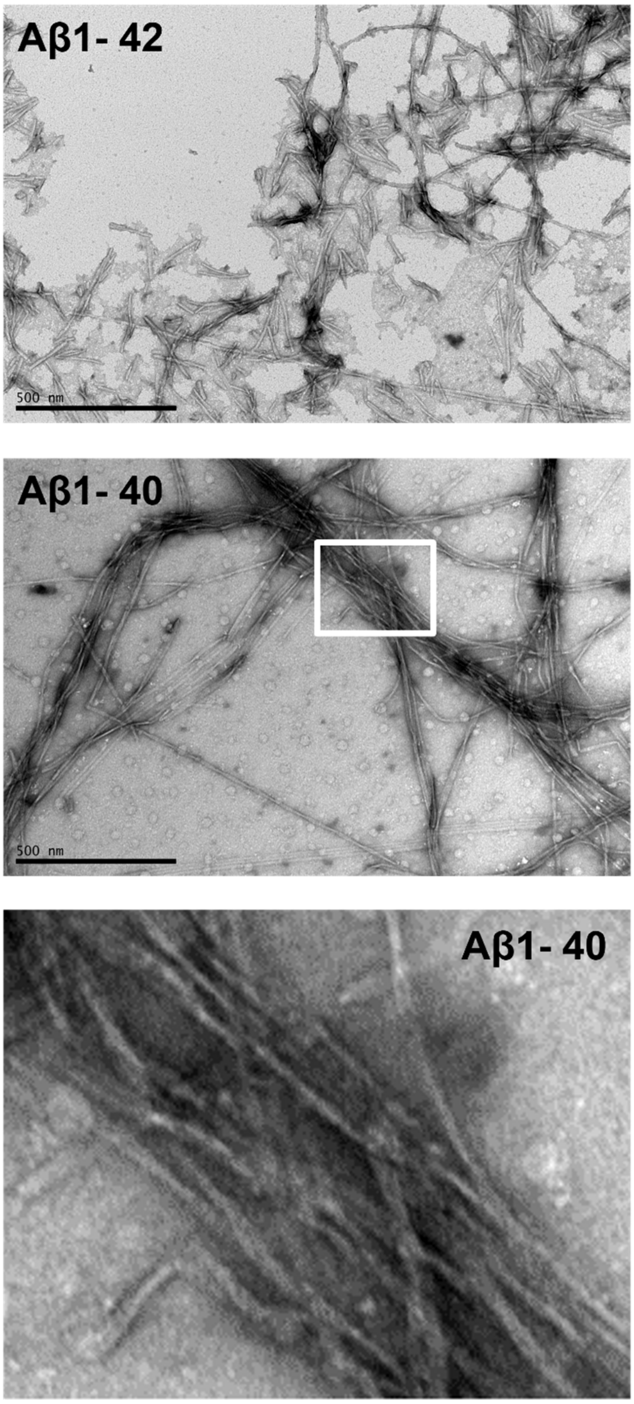


**Figure S1**: Negative stain TEM images of recombinant Aβ1-42 and Aβ1-40 fibrils fibrillated at 10 μM in PBS buffer pH 7.4 at 37°C without shaking. (A) TEM image of Aβ1-42 fibrils at 3-hour timepoint. The fibrils are short and unbundled. (B) TEM image of Aβ1-40 fibrils at 10-hour timepoint. The fibrils are very long and form bundles of intertwined fibrils (marked with a white box). (C) Zoomed view of the bundled Aβ1-40 fibrils. Scale bars are at 500 nm.

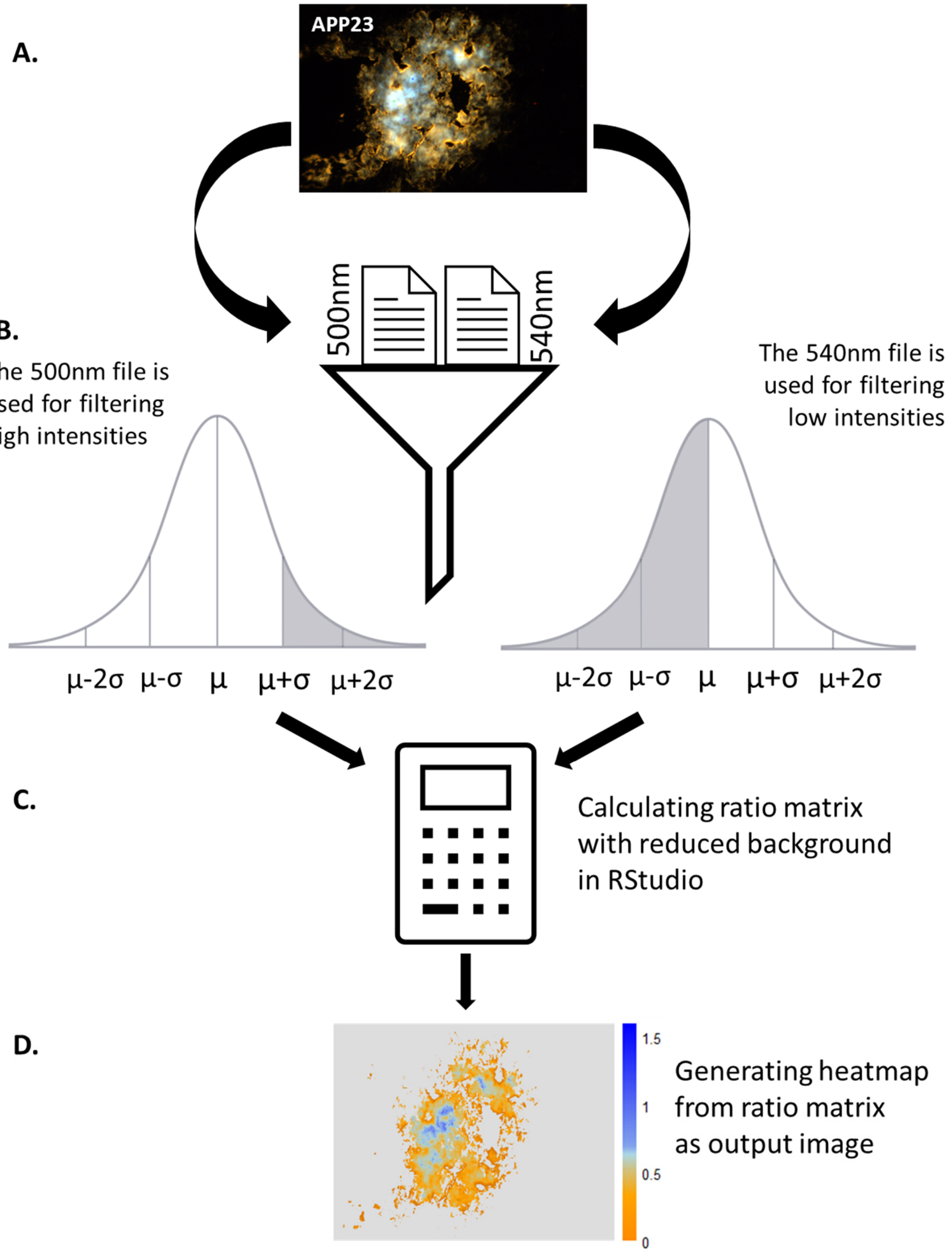


**Figure S2**: A schematic overview of image filtration using a relative filter setting in RStudio. (A) Overview of the raw image. (B) Text files at different wavelengths i.e., at 500 nm, 540 nm and 588 nm is extracted from the hyperspectral microscope and used to apply relative filter setting to remove unwanted high and low intensities from the raw image. (C) Ratio matrix is calculated using the filtered text files at desired wavelengths i.e., at $I_{500}/I_{540}$nm or $I_{500}/I_{588}$nm in RStudio. (D) A heatmap is generated from the ratio matrix as an output image without unwanted signal.

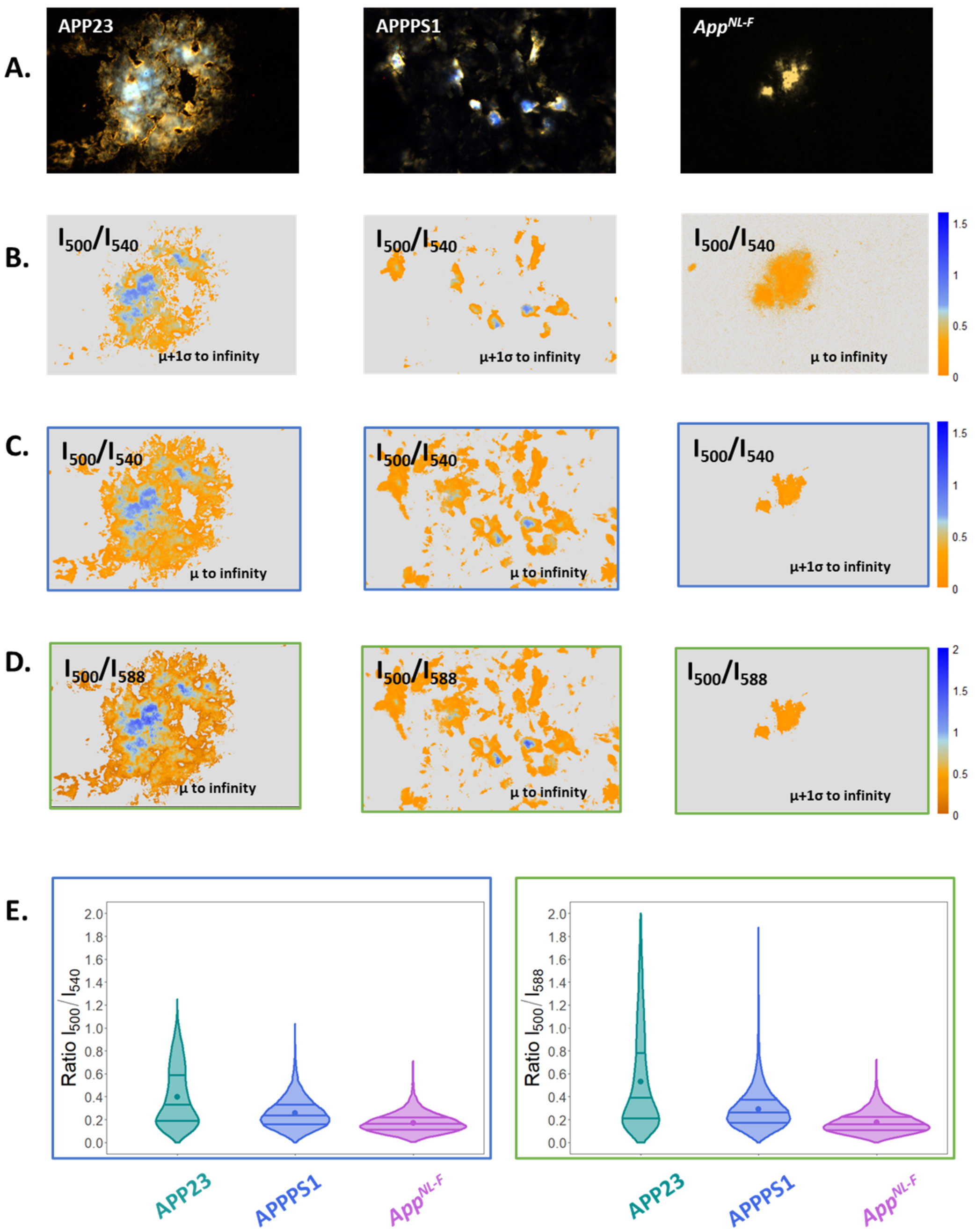


***Figure S3***: *(A) Overview of hyperspectral images of qFTAA and hFTAA stained amyloid plaques from APP23, APPPS1 and App$^{NL\text{-}F}$ mouse before applying the relative filter settings to the images. (B) Overview of the same plaques after applying the filter setting that is suboptimal for these genotypes i.e., APP23, APPPS1 and App$^{NL\text{-}F}$. (C) Overview of the plaques with the best relative filter setting at the intensity ratio of $I_{500}/I_{540}$ nm encircled with the blue box. For APP23 and APPPS1 mice lower limit of filter setting is μ, meanwhile for App$^{NL\text{-}F}$ mouse is μ+1σ. The upper limit is infinity for all the mouse models. (D) Overview of the plaques with the best relative filter setting at the intensity ratio of $I_{500}/I_{588}$ nm encircled with the green box. The filter setting for all the genotypes is the same as at the intensity ratio $I_{500}/I_{540}$ nm. (E) The violin plots represent the ratiometric analysis of the different genotypes at intensities $I_{500}/I_{540}$ nm and $I_{500}/I_{588}$ nm as before, but now considering all the pixels from images instead of ROI selection. The dots in each violin plot represent the mean value of the corresponding genotype. The straight lines from the bottom of the violin plot represent the 1$^{st}$ quartile (25% data below this line), 2$^{nd}$ quartile or median (50% data below this line) and 3$^{rd}$ quartile (75% data below this line) respectively.*

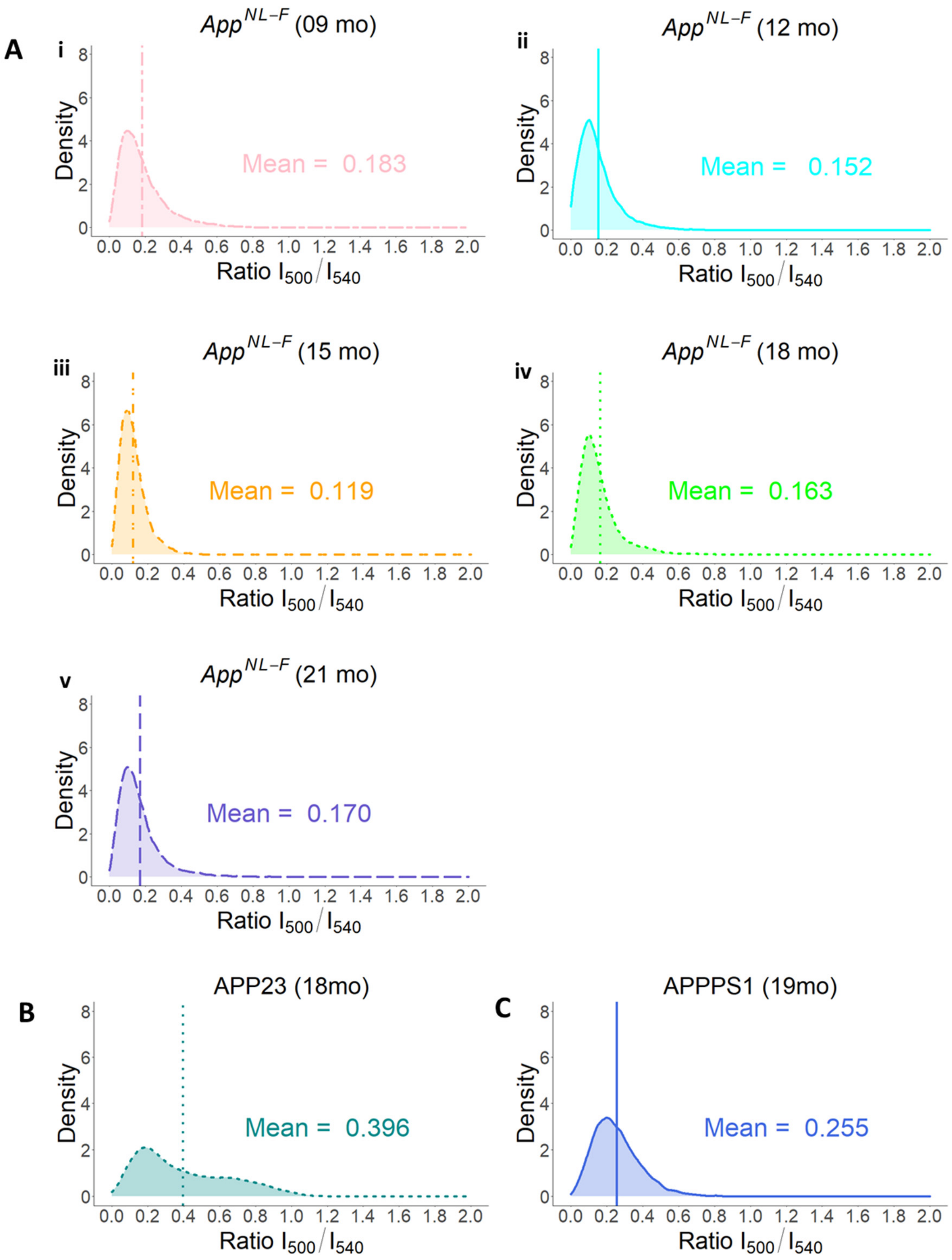


**Figure S4**: (A) Individual density plots of *App*$^{NL\text{-}F}$ mice at different ages: i) 9 months old mouse group with a mean intensity ratio vale 0.183 ii) 12 months old mouse group with a mean intensity ratio value of 0.152 iii) 15 months old mouse group with a mean intensity ratio vale 0.119 iv) 18 months old group mouse with a mean intensity ratio vale 0.163 v) 21 months old mouse group with a mean intensity ratio vale 0.163 (B) 18 months old APP23 mouse with a mean intensity ratio value of 0.396 (C) 19 months old APPPS1 mouse with a mean intensity ratio value of 0.255.
